# Ecological and evolutionary dynamics of cell-virus-virophage systems

**DOI:** 10.1101/2023.02.07.527428

**Authors:** Jose Gabriel Nino Barreat, Aris Katzourakis

## Abstract

Microbial eukaryotes can be infected by giant viruses, which can be infected by virophages. Virophages are parasites of the virus transcription machinery and can interfere with virus replication, resulting in a benefit to the eukaryotic host population. Surprisingly, virophages can integrate into the genomes of their cell or virus hosts, and have been shown to reactivate during coinfection. This raises interesting questions about the role of integration in the dynamics of cell-virus-virophage systems. Here, we use mathematical models and computational simulations to understand the effect of virophage integration on populations of cells and viruses. We also investigate programmed-cell death (PCD) and multicellularity as potential antiviral defence strategies used by cells. We found that virophages which enter the cell independently of the host virus, such as Mavirus, are expected to integrate commonly into the genomes of their cell hosts. In contrast, we show that virophages like Sputnik which form a complex with the giant virus, should rarely be found integrated in cell genomes. Alternatively, we found that Sputnik virophages can stably persist integrated in the virus population, as long as they do not completely inhibit virus replication. We also show that increasing virophage inhibition can stabilise oscillatory dynamics, which may explain the long-term persistence of viruses and virophages in the environment. Our results demonstrate that inhibition by virophages, PCD and multicellularity are effective antiviral strategies that may act in synergy against viral infection in microbial species.

## Introduction

Giant viruses are large dsDNA viruses from multiple families that belong to the clade of Nucleocytoplasmic Large DNA Viruses (NCLDVs) (1). They are known to infect a wide variety of eukaryotes including algae, protozoans and vertebrates (1), and some are amongst the largest viruses known in terms of genome size and physical dimensions; which can be easily seen under the light microscope (2–4). The striking sizes of giant viruses, which rival those of bacteria, are believed to have evolved independently from smaller viral ancestors via extensive horizontal gene transfer (5,6). A distinctive feature in the infectious cycle of giant viruses is the formation of electron-dense areas in the cell cytoplasm where viral transcription and assembly take place (7). These viral factories concentrate significant cell resources and are the sites where virally-encoded components of the transcription and translation machineries are expressed (8). The newly assembled viruses exit by lysis or budding, which results in the death of the infected cell (9).

Virophages are viral parasites of giant viruses which use their viral factories for replication. The first virophage to be described was Sputnik, found in association with *Acanthamoeba polyphaga mimivirus* (10). Sputnik could not replicate in the absence of its host virus and when grown together, caused the appearance of abortive and misassembled virus particles (10). Another class of virophages was then discovered in the flagellate *Cafeteria burkhardae* (formerly *Cafeteria roenbergensis*) infected by *Cafeteria roenbergensis virus* (CroV) (11). Similar to Sputnik, Mavirus required coinfection of the cell with CroV and interfered with CroV replication (11). The Guarani virophage was also shown to have this inhibitory effect on the replication of its mimivirus host (12). In general, it seems that virophages benefit their eukaryotic hosts by lowering the giant virus progeny and increasing the survival of the cell population (12,13). However, this might not be universal as the Zamilon virophage does not seem to have a significant impact on its host virus (14), and so it is a “neutral” virophage.

Virophages can have different biological properties which are illustrated by Mavirus and Sputnik. Mavirus can infect the cell independently of its host virus via receptor-mediated endocytosis (11), while Sputnik enters the cell forming a complex with its host mimivirus by attaching to glycosylated protein fibres of the outer capsid (15,16). Virophages can also integrate using different enzymes: a tyrosine recombinase in Sputnik or a rve-integrase in Mavirus (17). Interestingly, only Mavirus-like virophages have been found in the genomes of their eukaryotic hosts (13,18). Sputnik-like virophages have been found integrated in the genomes of the giant virus but not in the eukaryotic host, despite attempts to find integrated virophages in several *Acanthamoeba* species (19,20). In the flagellate *Cafeteria*, reactivation of integrated Mavirus virophages works as an inducible defence system against CroV infection (13). Although the integrated virophage does not seem to inhibit the virus host and infection leads to lysis, exogenous virophages are produced which can interfere with viral replication in new rounds of coinfection, protecting the wider host population (13).

Giant viruses and virophages are emerging as important players in the function of aquatic ecosystems, as evidenced by their sheer abundance and diversity across globally distributed metagenomes (21–24). They also seem to be ancient components of the eukaryotic virome (25– 27), and could thus be implicated in the evolution of antiviral strategies used by the early eukaryotes. Here we use mathematical models and computational simulations to study the ecological and evolutionary dynamics of this tripartite parasitic system. Importantly, we consider the role of virophage integration on the system dynamics which had not been studied in previous theoretical work. Our models explain why different virophages follow different integration strategies, they show that virus inhibition by virophages can stabilise the dynamics of the system and that inhibitory virophages, multicellularity and programmed-cell death may be used as effective antiviral strategies. Taken together, our findings shed light on various aspects of the ecology and evolutionary genomics of virophages and their hosts, and provide a framework of testable hypotheses to inspire future experimental work.

## Results

### ODE models

We first examined the system dynamics in the absence of virophages (Fig. 1, upper row). In both models, cells and viruses engaged in oscillatory predator-prey interactions. This oscillatory behaviour has also been observed in other models where cells are killed by lytic viruses (24,28,29). In contrast to Lotka-Volterra predator-prey models where trajectories converge to different orbits depending on initial conditions (neutral oscillations), our models show trajectories that converge to a limit cycle attractor in 2 dimensions (a stable oscillatory regime). This is similar to the behaviour observed in Holling-Tanner predator-prey models where population growth is also controlled by a logistic term (30,31). The appearance of an attractor is a desirable property since the system outcome is robust to small perturbations in the initial conditions (30).

**Fig 1.**
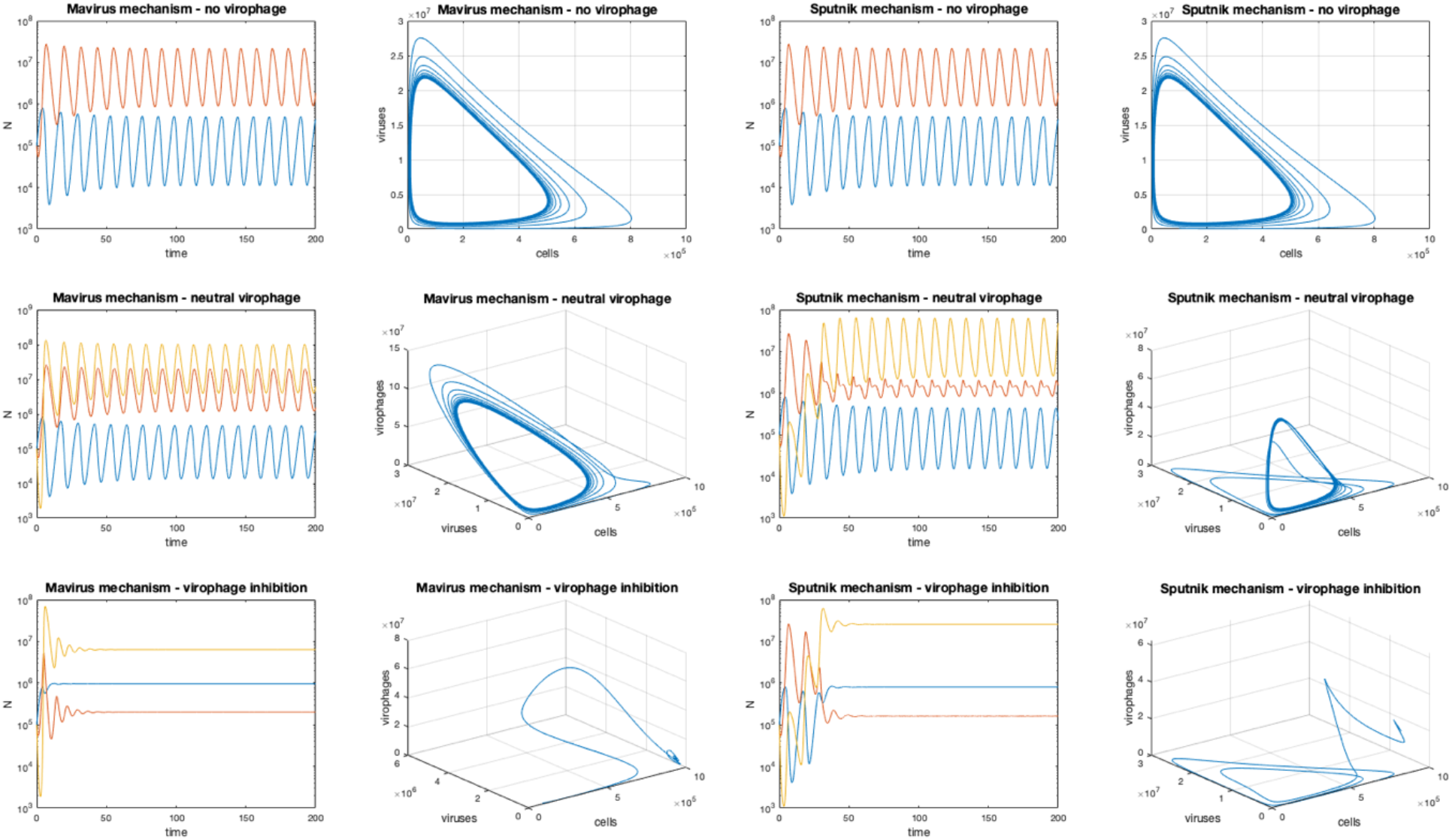
Dynamics of the Mavirus and Sputnik 1a models in the absence of virophage integration (baseline scenario). In the time-course diagrams, the blue line represents the population of cells, the orange represents viruses and the yellow line represents virophages. **Upper row:** cell-virus interactions produce Holling-Tanner oscillations, trajectories converge to a 2-dimensional limit cycle in state space. **Middle row:** when neutral virophages are added, the dynamics are still oscillatory, trajectories converge to a 3-dimensional limit cycle. **Bottom row:** total inhibition by virophages stabilises the system dynamics, trajectories now converge to a point equilibrium. Parameters for model 1: **α**_**1**_ = 1, **α**_**2**_ = 0.9, **β**_**1**_ = 10^−7^, **β**_**2**_= 10^−6^, **γ**_**1**_ = 1.2, **γ**_**2**_ = 2.6, **λ** = 0, **φ**_**1**_ = 100, **φ**_**2**_ = 1000, **f** = {1, 0}, **K** = 10^6^, **r**_**1**_ = 1, **r**_**2**_ = 0.8, **p** = 0. Parameters for model 2 are the same except for **β**_**2**_ = 10^−7^ and **k** = 8·10^−7^. Initial conditions: **C**_**x**,**0**_ = 10^5^, **G**_**0**_ = 10^5^, **V**_**0**_ = 10^5^.

When we added neutral virophages to the system (which do not inhibit virus replication), all three populations showed a stable oscillatory regime (Fig. 1, middle row). This can be seen in the appearance in state-space of a 3-dimensional limit cycle attractor. Interestingly, when we set the inhibition parameter equal to zero (**f = 0**, virophages completely inhibit virus replication), the trajectories converge to a point equilibrium where the populations of cells, viruses and virophages remain unchanged. This is an example of a Hopf bifurcation, where a fixed-point equilibrium gives rise to a limit cycle attractor (30,31), in this case as a function of the inhibition parameter (**f**).

### The effect of virophage integration into the cell genome

Inclusion of integrating virophages affected dynamics of the system differently depending on whether virophages followed the Mavirus or Sputnik infection mechanisms. When virophages are neutral and the rate of integration is low (**f** = 1, ***λ*** = 10^−9^), we observed the appearance of cells with an integrated virophage which oscillated together with naïve cells, virophages and viruses in the Mavirus model (Fig. 2, upper row). In contrast, we did not observe cells with an integrated virophage emerging in the model for Sputnik under these conditions (Fig. 2, upper row). If we then increased the integration rate (**f** = 1, ***λ*** = 10^−4^), cells with an integrated virophage were observed in the Sputnik model, while these cells replaced the naïve cell population in the Mavirus model (Fig. 2, upper middle row). Keeping the integration rate constant but in presence of moderate virophage inhibition (**f** = 0.6, ***λ*** = 10^−4^), led to the appearance of a point equilibrium with an “inward spiral” pattern in state-space (Fig. 2, lower middle row). In the Mavirus model, naïve cells were replaced by cells with an integrated virophage, while in the Sputnik model both cell populations were present. Under total virophage inhibition and low integration rate (**f** = 0, ***λ*** = 5×10^−8^), both models also showed damped oscillations converging to a point equilibrium, with replacement of naïve cells in the Mavirus model and stable coexistence in the Sputnik model (Fig. 2, lower panel).

**Fig 2.**
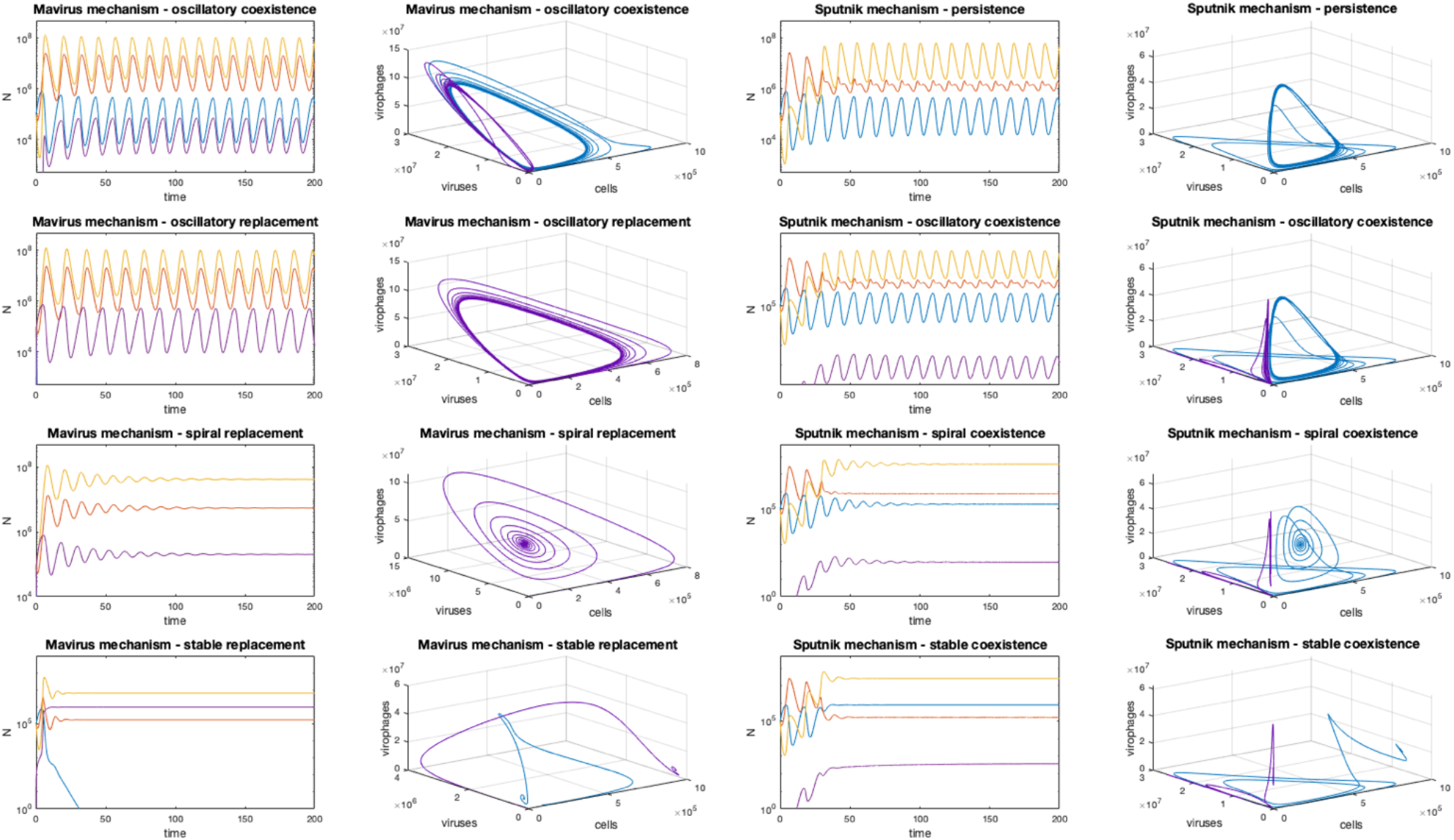
Dynamics of the Mavirus and Sputnik 1a models in the presence of virophage integration. In the time-course diagrams, the blue line represents the population of naïve cells, the purple line represents cells with an integrated virophage, the orange represents viruses and the yellow line represents virophages. **Upper row:** in the presence of neutral virophages and low rates of integration (**f** = 1, ***λ*** = 10^−9^), we observe oscillatory coexistence of all populations in the Mavirus model while cells with an integrated virophage cannot establish in the Sputnik mechanism. The Mavirus trajectories converge to a 4-dimensional limit cycle in state-space (shown as two nested trajectories), while the Sputnik trajectories converge to a 3-dimensional limit cycle. **Upper middle row:** in the presence of neutral virophages and higher integration rates (**f** = 1, ***λ*** = 10^−4^), cells with an integration become fixed in the population in the Mavirus model (3-dimensional limit cycle), while coexistence of all four populations was observed in the Sputnik model (4-dimensional limit cycle). **Bottom middle row:** in the presence of moderate inhibition by virophages (**f** = 0.6, ***λ*** = 10^−4^), damping-wave oscillations are observed in both systems but fixation occurs in the Mavirus mechanism while coexistence occurs with Sputnik. Populations converge to a point equilibrium with spiral trajectories. **Bottom row:** total inhibition leads to stabilisation in both systems (**f** = 0, ***λ*** = 5e-8), with fixation of cells with an integrated virophage for Mavirus and coexistence for Sputnik. Parameters for model 1: **α**_**1**_ = 1, **α**_**2**_ = 0.9, **β**_**1**_ = 10^−7^, **β**_**2**_ = 10^−6^, **γ**_**1**_ = 1.2, **γ**_**2**_ = 2.6, **φ**_**1**_ = 100, **φ**_**2**_ = 1000, **K** = 10^6^, **r**_**1**_ = 1, **r**_**2**_ = 0.8, **p** = 0. Parameters for model 2 are the same except for **β**_**2**_ = 10^−7^ and **k** = 8·10^−7^. Initial conditions: **C**_**x**,**0**_ = 10^5^, **G**_**0**_ = 10^5^, **V**_**0**_ = 10^5^.

We examined these effects on the system outcomes more systematically by sampling the inhibition parameter and the integration rate over multiple orders of magnitude (Fig. 3). For both models, we observed that the main parameter determining system outcome is the integration rate, although it does interact to an extent with the inhibition parameter. In the Mavirus model, moderate rates of integration led to coexistence of naïve cells with cells carrying an integrated virophage (green region), while higher integration rates led to replacement of naïve cells with cells carrying an integrated virophage (red region). In contrast, the outcomes of the Sputnik model were dominated by naïve cells alone (blue region), while coexistence of cell populations was only observed at high rates of integration; higher than those required to observe replacement in the Mavirus model (green region).

**Fig 3.**
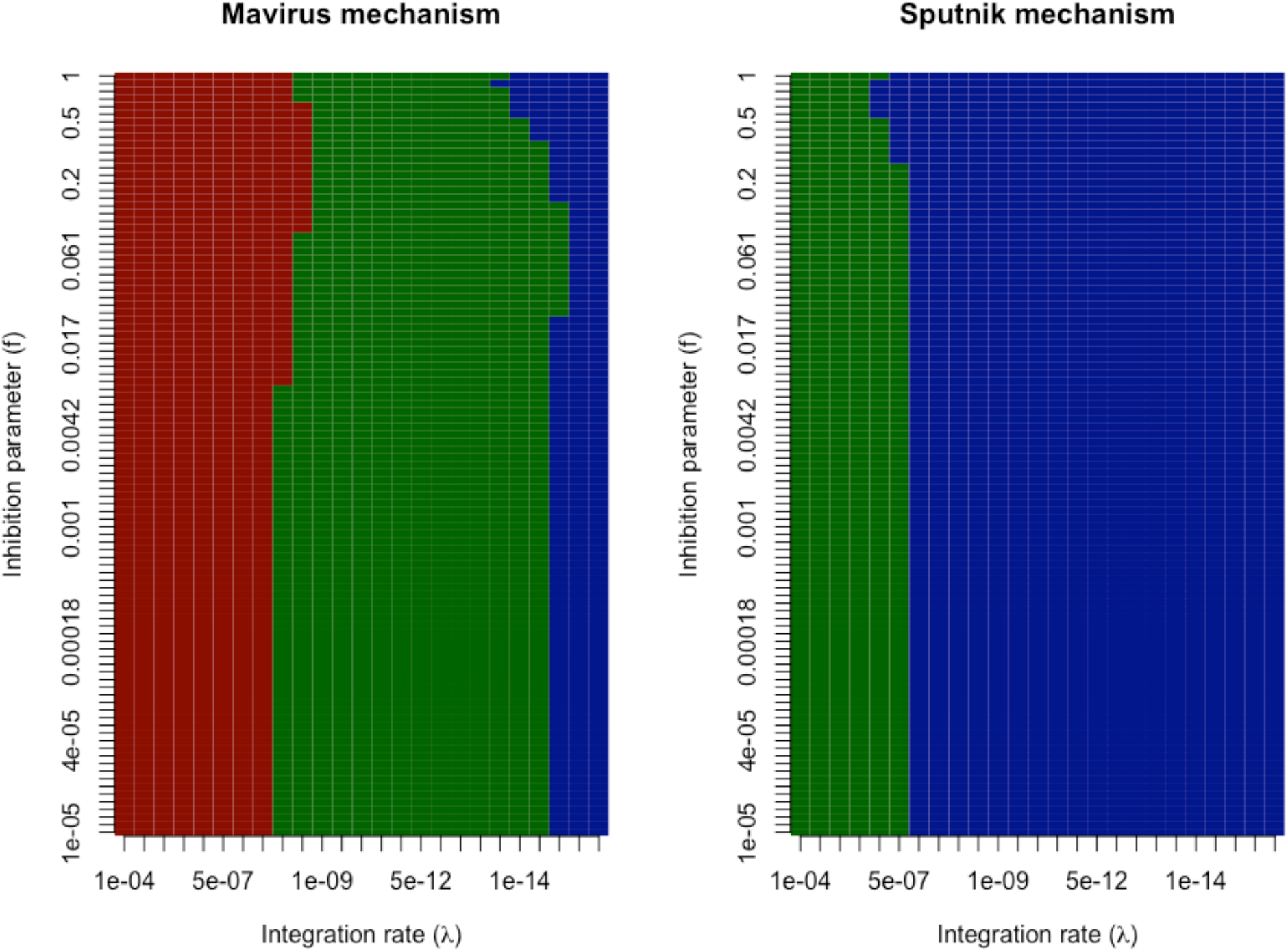
System outcomes depending on the virophage infection mechanism. We explored the outcome of the Mavirus and Sputnik models by varying the integration rates (***λ***) and the inhibition parameter (**f**) and mapping the dynamical outcomes. Mavirus **model (left)**: we observed three possible outcomes, fixation of cells with an integrated virophage (red), coexistence of cells with and without an integrated virophage (green) and persistence of cells without a virophage integration (blue). Sputnik **model (right)**: two outcomes were observed for the same parameter ranges, coexistence of cells with and without and integrated virophage (green) and persistence of naïve cells (blue). Parameters for model 1: **α**_**1**_ = 1, **α**_**2**_ = 0.9, **β**_**1**_ = 10^− 7^, **β**_**2**_ = 10^−6^, **γ**_**1**_ = 1.2, **γ**_**2**_ = 2.6, **φ**_**1**_ = 100, **φ**_**2**_ = 1000, **K** = 10^6^, **r**_**1**_ = 1, **r**_**2**_ = 0.8, **p** = 0. Parameters for model 2 are the same except for **β**_**2**_ = 10^−7^ and **k** = 8·10^−7^. Initial conditions: **C**_**x**,**0**_ = 10^5^, **G**_**0**_ = 10^5^, **V**_**0**_ = 10^5^.

### The effect of virus-induced PCD

The introduction of PCD triggered by viral infection had a similar stabilising effect on the system dynamics as observed for virophage inhibition (S1 Fig.). We considered this case by setting the PCD parameter to be greater than zero (**p** > 0). Compared to a model where Mavirus shows oscillatory replacement (**f** = 1, ***λ*** = 10^−7^, **p** = 0), inclusion of the PCD parameter (**p** = 6) led to stabilisation and convergence to a fixed-point equilibrium (S1 Fig., upper row). If the PCD was increased (**p** = 8), the cells reached a fixed-point equilibrium with an integration polymorphism while viruses and virophages went extinct (S1 Fig., upper row). Compared to an oscillatory persistence regime for the Sputnik model (**f** = 1, ***λ*** = 10^−7^, **p** = 0), setting **p** = 6 also led to stabilisation of the dynamics and convergence to a point equilibrium but virophages went extinct (S1 Fig., lower row). Increasing the PCD parameter to 8 led to the extinction of viruses and virophages, while naïve cells grew to carrying capacity (S1 Fig., lower row). The extinction of viruses and virophages occurred in both models with **p** ;: 6.6. In the Mavirus model, the time to extinction of viruses and virophages decreases as the PCD parameter is increased (S1 Fig., third column). The time to extinction of virophages was independent of increases in the PCD parameter in the Sputnik model, while the time to extinction of viruses also decreased as the PCD parameter was increased.

### Integration of Sputnik into the virus genome

We explored integration of Sputnik into the host virus genome since it has been observed experimentally (19). In the absence of integrating virophages (***λ*\*** = 0), the system showed the same oscillatory dynamics with neutral virophages and damping oscillations under virophage inhibition, as described for the previous two models (Fig. 4, upper row). An integration frequency of 1/10,000 (***λ*\*** = 10^−4^) led to the appearance of viruses with an integrated virophage under no to moderate inhibition (**f** = {1, 0.7}; Fig. 4, middle row). However, in the presence of total virophage inhibition (**f** = 0), viruses with an integrated virophage did not establish in the population (Fig. 4, middle row). This makes intuitive sense since a virus carrying a totally inhibitory virophage would not be able to replicate. Interestingly, when 1/10 viruses produced during a mixed infection carried an integrated virophage (***λ*\*** = 10^−1^), these viruses outcompeted the ones without an integration if the virophages were neutral or moderately inhibitory (**f** = {1, 0.7}), but they were not observed if inhibition was total (Fig. 4, lower row).

**Fig 4.**
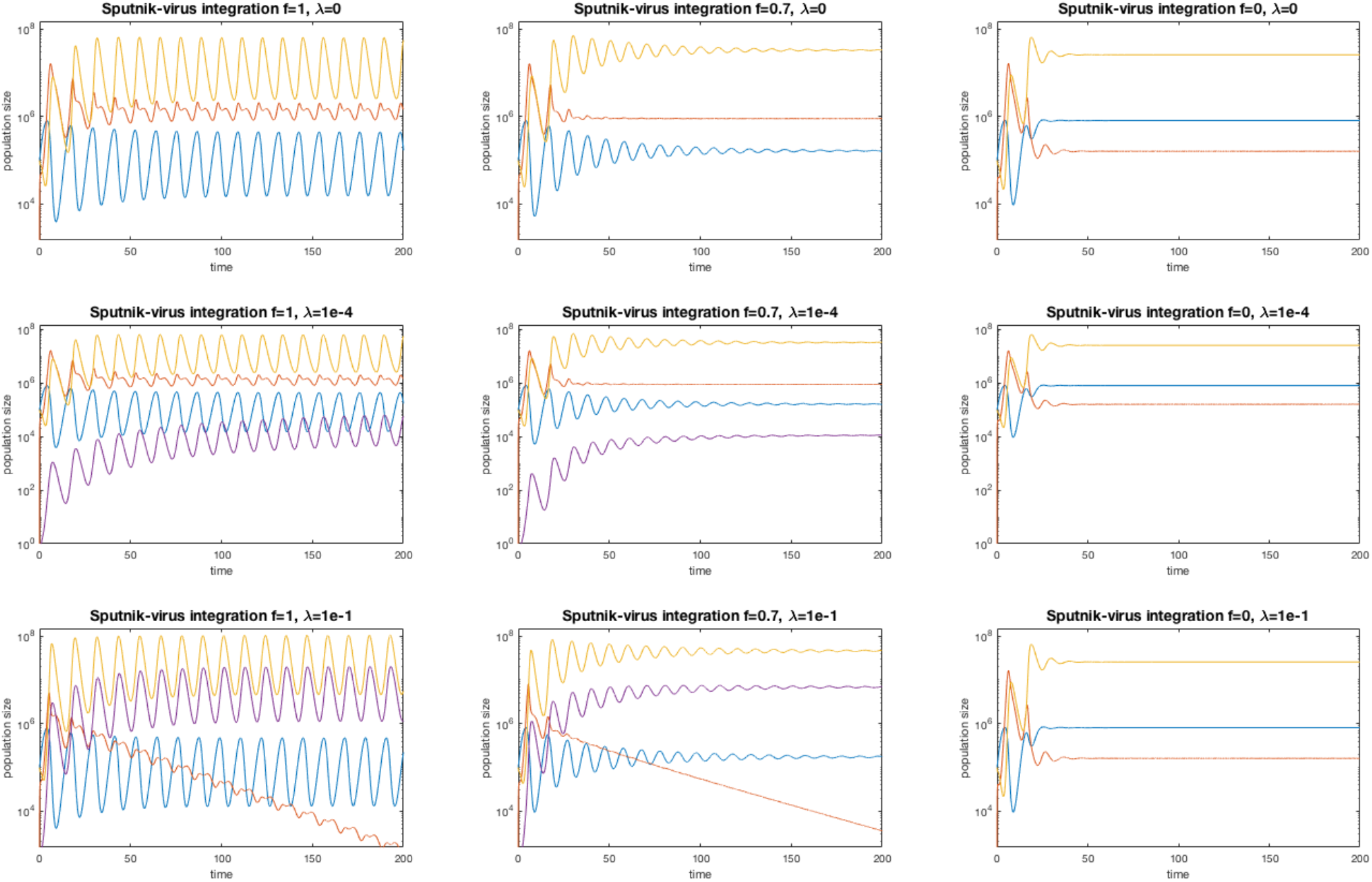
Dynamics of the Sputnik 1b model with integration into the host virus genome. In the time-course diagrams, the blue line represents the population of cells, the orange represents viruses, the purple line represents viruses with an integrated virophage and the yellow line represents virophages. Virophage inhibition leads to stabilisation of the dynamics. **Upper row:** in the absence of integration (***λ*\*** = 0), the cell, virus and virophage populations coexist in time. **Middle row:** in the presence of moderate integration (***λ*\*** = 10^−4^), viruses with an integrated virophage can establish in the population and are maintained as a polymorphism (**f** = {1, 0.7}), unless there is total virophage inhibition (**f** = 0). **Bottom row:** in the presence of high integration (***λ*\*** = 0.1), the original virus population is replaced by viruses carrying an integrated virophage (**f** = {1, 0.7}), except when inhibition is total (**f** = 0).

### Agent-based model

We developed an agent-based model (ABM) in 3-dimensions to understand the impact of spatial effects in a stochastic simulation of cell-virus-virophage interactions. In particular, we were interested in understanding the effect of multicellularity on the system dynamics and how it interacted with virophage inhibition and PCD. For this, we considered 15 different scenarios in which populations of organisms with different cell numbers (single-, 2-, 4-, 8- and 16-celled organisms) were exposed to neutral virophages, inhibitory virophages or PCD. Our ABM considered a population of non-dividing cells in order to reduce the computational complexity of the simulations, and thus centres on a single-epidemic wave.

### The effect of virophage inhibition, multicellularity and PCD

In the simulations we can see how the cell population was infected by viruses producing clouds of infection and bursts of virophages whenever they happened to coinfect a cell (S1-S15 Videos, DOI: <u>10.6084/m9.figshare.19412066</u>). An example time-course gives an idea of the underlying dynamics (S1-S3 Figs.). In a population of single-celled organisms, the neutral virophage treatment produced the largest waves of viruses and the lowest numbers of cell survivors. Both virophage inhibition and PCD appeared to have a protective effect on the cell populations. Increasing the number of cells per organism (which introduces spatial clustering of cells), also had a protective effect since the waves for viruses were lower and it increased the number of cell survivors compared to the single-celled case.

To assess these effects systematically, we carried out 100 replicates of the stochastic simulations per treatment and then analysed them using Generalised Additive Models (GAMs, DOI: <u>10.6084/m9.figshare.19412468</u>). Compared to a population of single-celled organisms, increasing the cell number led to a general decrease in the maximum amplitude of the virus wave up to 8-celled organisms, although an increase was then observed from the 8- to 16-celled populations (Fig. 5). The treatments with virophage inhibition had lower maximum amplitudes of the virus wave compared to neutral virophages, and the most effective treatment was PCD (Fig. 5). Multicellularity also lowered the time to virus extinction across treatments, and the protective effects of virophage inhibition and PCD were also observed (Fig. 6). Finally, the number of surviving cells was highest for the PCD treatment, followed by the treatment with inhibitory virophages, and increased as a function of the number of cells per organism (Fig. 7).

**Fig 5.**
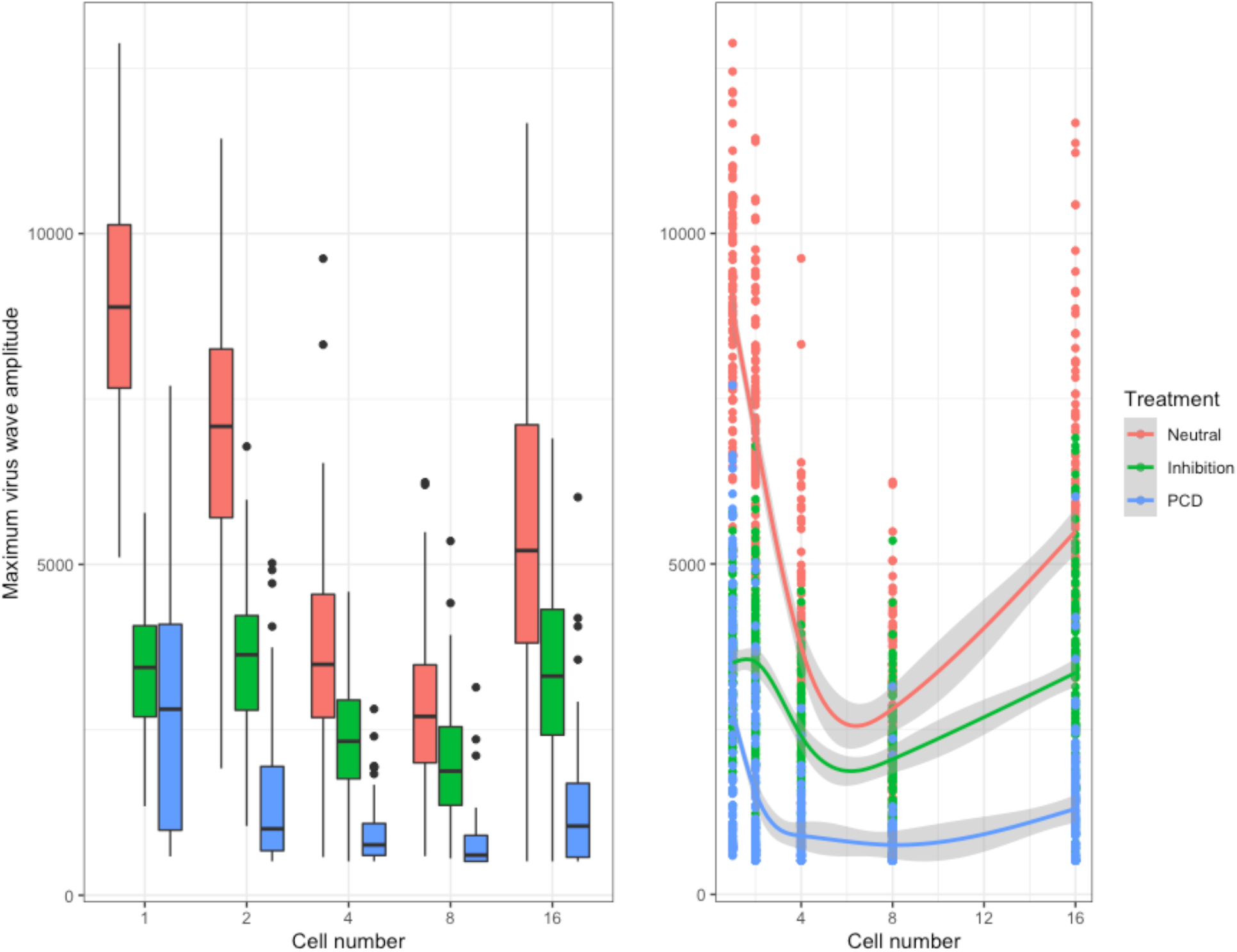
Maximum amplitude of the virus wave. The maximum amplitude of the virus wave decreases as a function of the number of cells and the treatment in the simulations. Boxplots of the resulting distributions are shown to the left and the fitted GAMs are shown to the right. However, note the increase in the maximum wave amplitude across all treatments from 8- to 16-celled organisms. The three intercepts are significantly different and the smoothing terms statistically significant to the level of p < 10^−15^. The normalised root mean square error (RMSE) is λ% and the deviance explained is 72%.

**Fig 6.**
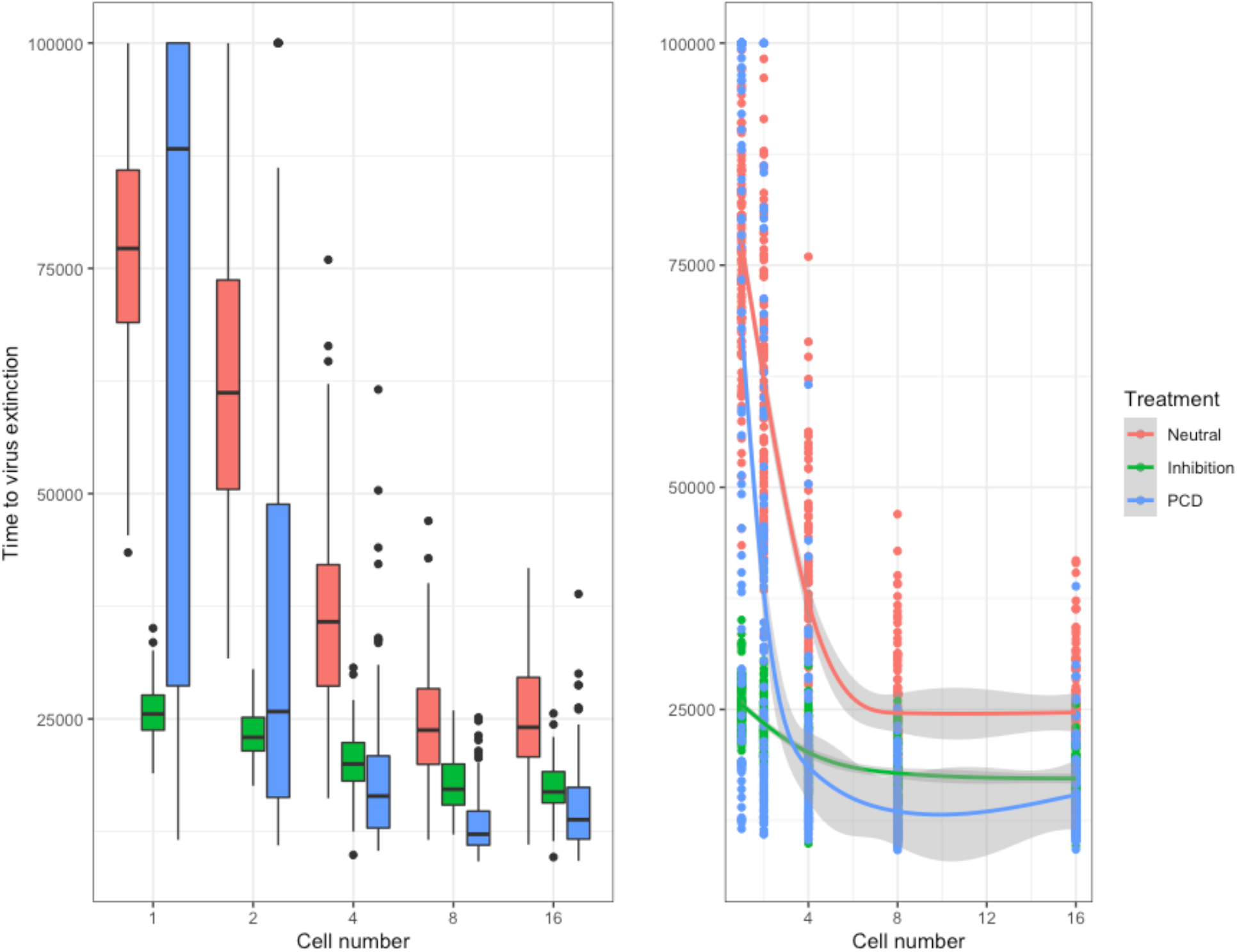
Time to virus extinction. The time to virus extinction also decreases as a function of the cell number and treatment in the simulations. Boxplots of the resulting distributions are shown to the left and the fitted GAMs are shown to the right. For single-celled organisms, virophage inhibition was more effective than PCD, since it led to faster extinction of viruses. The treatments with neutral and inhibitory virophages appear to level-off with increasing cell number, while there was an increase in the time to virus extinction in the PCD treatment from 8- to 16-celled organisms (despite the errors being larger). The three intercepts are significantly different and the smoothing terms statistically significant for the neutral virophages and PCD to the level of p < 10^−15^. The smoothing term for the treatment with inhibitory virophages was significant to the level of p < 10^−5^. The normalised root mean squared error (RMSE) is 14.6% and the deviance explained is 69.3%.

**Fig 7.**
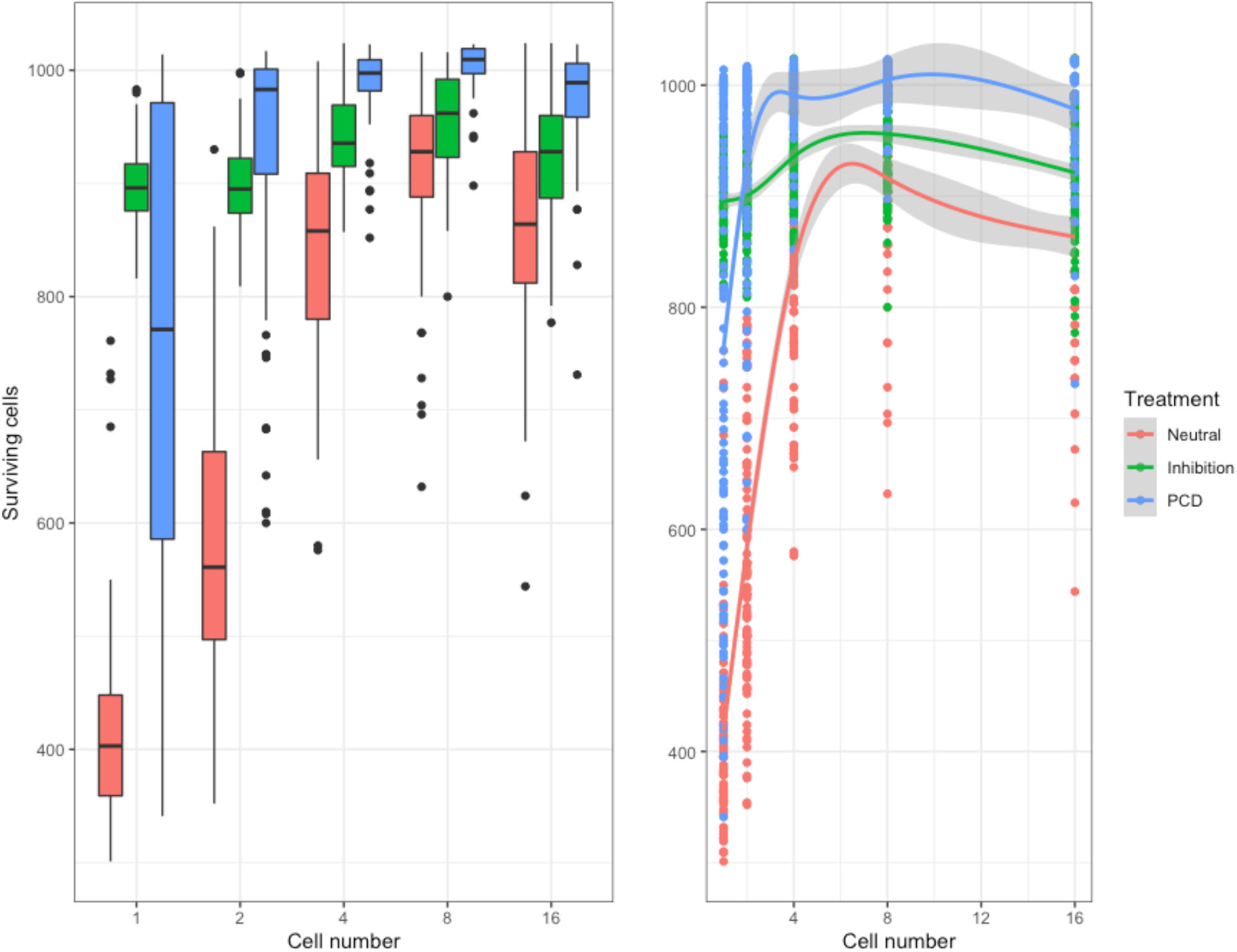
Number of surviving cells. The number of surviving cells at the end of the simulations increased as a function of the cell number and the treatment. Boxplots of the resulting distributions are shown to the left and the fitted GAMs are shown to the right. The highest levels of survival were observed for the PCD treatment, followed by virophage inhibition. The three intercepts are significantly different and the smoothing terms statistically significant for the neutral virophages and PCD to the level of p < 10^−15^. The smoothing term for the treatment with inhibitory virophages was significant to the level of p < 10^−5^. The normalised root mean squared error (RMSE) is 0.7% and the deviance explained is 76.8%.

**Fig 8.**
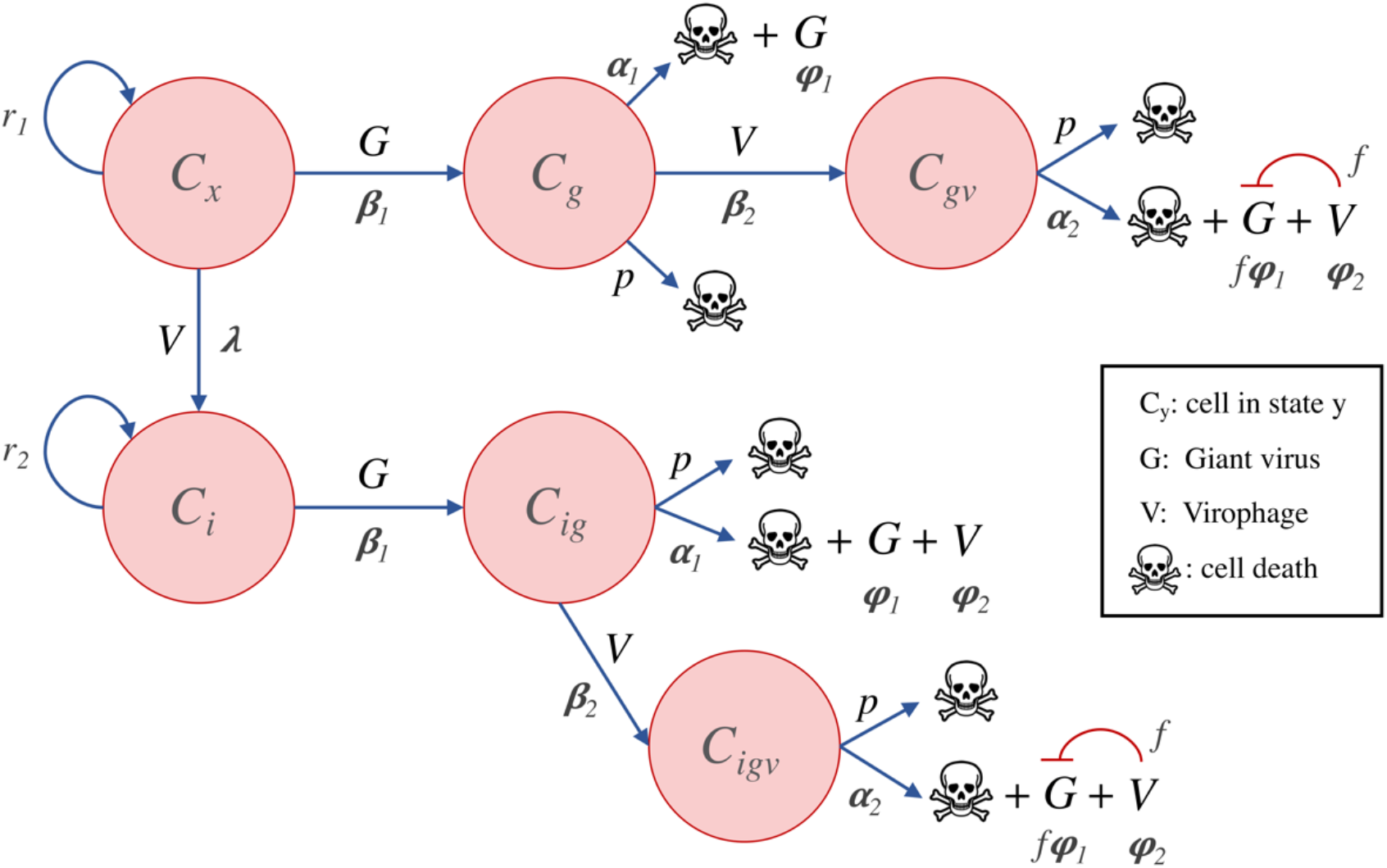
Model of virophage integration into the cell genome following the Mavirus mechanism. Cells with (**C**_**i**_) or without an integrated virophage (**C**_**x**_) can be infected by a virus. Infection of **C**_**x**_ cells by a virus results in lysis and release of viruses, while infection of **C**_**i**_ cells by a virus leads to virophage reactivation and release of viruses without inhibition. Cells infected by a virus (**C**_**g**_ or **C**_**ig**_) can also be infected by a virophage, which results in lysis, release of virophages and virus inhibition

## Discussion

Our findings reveal several key aspects of the natural history of virophages and the interactions with their viral and cellular hosts. We have shown that increasing the inhibition of virophages on virus replication can have a stabilising effect on the oscillatory dynamics. This is important since non-oscillatory stable systems are more robust to stochastic variations that can lead to collapse of the populations (32). A previous model that did not consider the populations of viruses and virophages explicitly, suggested that increasing inhibition would be destabilising, leading to oscillatory dynamics and making the populations prone to extinction (32). To reconciliate this with the observed abundance and persistence of virophages in nature (23,33), the existence of a viral reservoir or meta-population dynamics were proposed (32). However, our models show that the long-term stability of virophages and giant viruses in nature is possible without the need of external intervening factors, although it is likely that multiple factors may be at play.

Another important observation that emerged from our analyses is that integration of virophages into the cell or virus genomes is potentially determined by the virophage infection mechanism. Virophages that follow an independent-entry mechanism such as Mavirus would be expected to be more commonly found in the genomes of their eukaryotic hosts, while viruses that enter the cells as a complex would be harder to be found integrated in cells. These predictions are consistent with experimental observations: Mavirus and Mavirus-like virophages have been found in the genomes of *Cafeteria burkhardae* and *Bigelowiella natans* (13,18,34), while Sputnik and Sputnik-like virophages have only been found integrated in the genomes of their virus hosts (17,19,20). One important aspect about the natural history of the Sputnik system is that a virophage integration in a cell can only survive if it occurs together with an abortive viral infection, otherwise the cell is lysed and its descendants will not inherit the integration (17).

In addition, we observed that when Sputnik virophages integrated into a high proportion of viral progeny (>1/10,000), viruses with integrated virophages could persist stably in the population. High rates of integration into the viral genome could indicate that Sputnik has evolved a greater affinity for integration into the virus compared to the host cell, probably as a consequence of its unique infection mechanism. Indeed, tyrosine recombinases do seem capable of integrating into the host cell genome, as shown by a *Polinton*-like virus that was discovered integrated in the genome of the cryptophyte alga *Guillardia theta* (35). It would be interesting to test if changing the infection mechanisms of Mavirus and Sputnik leads to different integration dynamics as predicted by our models. For example, Mavirus and Sputnik may be genetically modified to use each other’s outer capsid proteins and host virus (i.e. Mavirus/APMV and Sputnik/CroV), and it could then be seen which integration pattern they follow.

Our analyses also indicate that virophage inhibition, virus-induced PCD and the transition to multicellularity can be effective antiviral strategies in microbial eukaryotes. PCD has been classically studied in the context of complex multicellular organisms, where is has important functions in development and homeostasis (36), but PCD also occurs in microbes (37). This gives rise to a paradox since the suicide of a single cell ends its reproductive potential. However, artificial life models have shown that kin selection is sufficient to explain the evolution of suicide in single-celled organisms (38). In *Escherichia coli*, cell-suicide has been shown to be advantageous in the presence of lytic phage T4*rII* even when relatedness is low: the benefit to the closest kin greatly exceeds the cost to the infected cell since it will die anyway (39). In the simulations, we observed that a multicellular population of organisms gains protection against viral infection compared to a population of single-celled organisms. Our explanation for this spatial effect is that as long as viruses decay over time, hosts that are more widely dispersed in space will be harder to reach. Therefore, it is more likely that the virus will decay if it has to traverse a longer diffusing distance, while in the single-celled case, it can use hosts as stepping-stones to travel through space more easily. Our results are consistent with previous modelling work that used a Markov process to show that cell-clustering and PCD are optimal strategies in the presence of high viral loads and imperfect immunity (40).

We have provided mechanistic insights into the interactions of virophages with their virus and cellular hosts, and at the same time explored antiviral defence strategies that are at the disposal of microbial eukaryotes. These results are relevant to the evolution of antiviral immunity in early eukaryotes and to the evolution of multicellularity. Indeed, there is evidence to support horizontal gene transfers between NCLDVs and proto-eukaryotes, which indicates these viral lineages already existed more than a billion years ago (25,27). Virophages are also believed to be an ancient group of viruses, given their diversity, abundance and association with multiple NCLDV host families (23,33,41). In light of this ancient arms-race, the early eukaryotes may have used inhibitory virophages to protect themselves against the onslaught from NCLDVs. The interaction with viruses may also have contributed to the evolution of PCD in eukaryotes and to the initial transition to multicellularity. In this regard, it would be interesting to test if simple colonial forms of unicellular eukaryotes can evolve in the presence of a lytic virus, as they can do when exposed to a predator (42,43). Other processes, such as the sharing of public goods, may have also played major roles in the evolution of multicellularity and PCD (38,44). Since they do not appear to be mutually exclusive, it seems likely that the interaction with viruses and the sharing of public goods may have acted together to favour the appearance of multicellular eukaryotes on multiple occasions. The extent to which viruses may be a driving or reinforcing factor during these transitions can be explored in future work.

## Materials and methods

We used two main approaches to study the dynamic interactions between cells, viruses and virophages in the presence of virophage integration. Our first approach was to design and study the qualitative behaviour of coupled systems of Ordinary Differential Equations (ODEs). These systems considered the different virophage infection mechanisms and integration into the cell or host virus genomes. The second approach was to use an Agent-based model (ABM) which allowed us to consider spatial effects and analyse the effect of multicellularity on the dynamics of the system. The codes and scripts developed to implement these models are available in GitHub (https://github.com/josegabrielnb/virophage-dynamics).

### ODE models

To study the impact different infection mechanisms and virophage integration, we developed three systems of coupled ODEs. Our systems of ODEs explicitly model the populations of cells, viruses and virophages, and assume that the system is well-mixed. The first system is based on the independent-entry mechanism of Mavirus and integration into the host cell genome. The second system describes the paired-entry mechanism of Sputnik (the virophage enters the cell in a complex with the virus), and integration into the cell genome. The third model is also based on the paired-entry mechanism but considers integration of Sputnik into the host virus genome, which has been observed experimentally (19).

### Mavirus model

The Mavirus model assumes that the virophage can enter the cell independently of its host virus (Fig 9). Naïve cells (**C**_**x**_) can be infected by viruses (**G**) or virophages (**V**). Cells infected by a virus (**C**_**g**_), die at a certain rate and produce viral progeny, or they can be infected by a virophage. Cells that are infected by a virus and a virophage (**C**_**gv**_), produce viral progeny and release virophages when the cell dies. The amount of viral progeny produced is determined by the inhibition parameter (**f**), where **f** = 0 is total inhibition and **f** = 1 is no inhibition (neutral virophages). In the absence of a host virus, a virophage may infect and integrate into the genome of a naïve cell (**C**_**i**_). When this cell encounters a giant virus (**C**_**ig**_), the integrated virophage is reactivated but does not lead to virus inhibition, which is in line with experimental observations (13). The virus-infected cell with a provirophage may also be infected by an exogenous virophage (**C**_**igv**_), which can have an inhibitory effect on viral replication. The model makes the simplifying assumption that virophages which infect a cell but which do not integrate are degraded. A similar assumption was made by Wodarz in model 1a (32). This was done to arrive at a simpler model and to allow a more direct comparison with the Sputnik mechanism..

**Fig 9.**
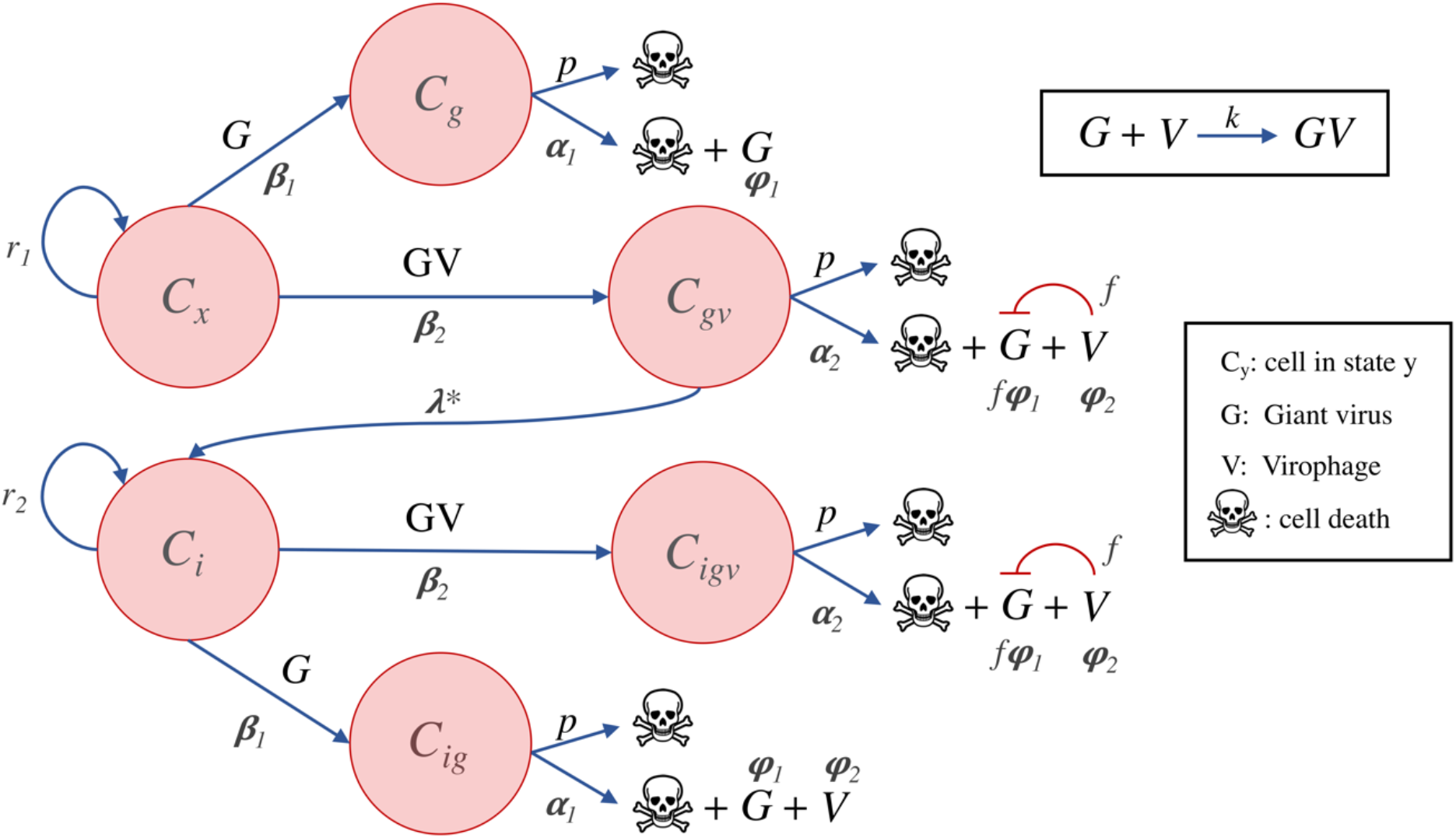
Model of virophage integration into the cell genome following the Sputnik mechanism. Cells with (**C**_**i**_) or without an integrated virophage (**C**_**x**_) can be infected by a virus or by a virus-virophage complex (**GV**). Infection of **C**_**x**_ cells by a virus results in lysis and virus release, while infection of **C**_**i**_ cells by a virus leads to virophage reactivation and release of viruses without inhibition. Infection of **C**_**x**_ or **C**_**i**_ cells by a complex, results in lysis, release of virophages and virus inhibition.

The Mavirus model is described by a system of 8 coupled ODEs and 14 parameters:

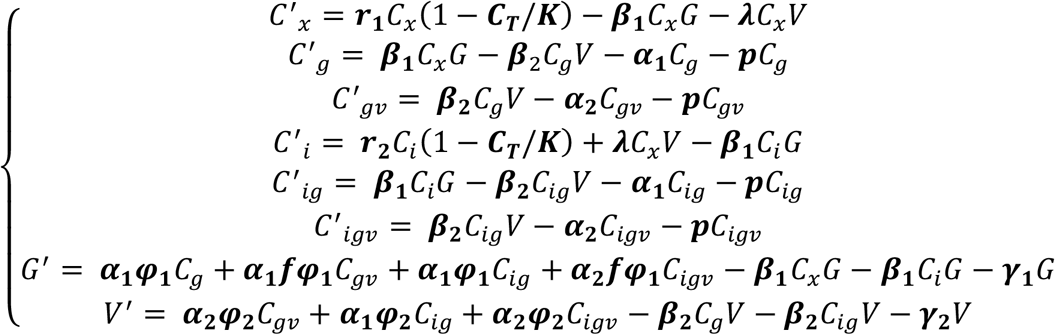

Where **r**_**1**_ and **r**_**2**_ are the intrinsic growth rates of naïve cells and cells carrying a provirophage, **K** is the carrying capacity of cells in the population, **β**_**1**_ and **β**_**2**_ are the infection rates of viruses and virophages, **α**_**1**_ and **α**_**2**_ are the death rates of cells infected by a virus or a virus and virophage, **φ 1** and **φ**_**2**_ are the virus and virophage burst sizes, **γ**_**1**_ and **γ**_**2**_ are the decay rates for viruses and virophages, **λ** is the integration rate of virophages into the cell genome, **f** is the degree of virophage inhibition and **p** is the PCD parameter. Cells can commit suicide in response to a viral infection if the PCD parameter (**p**) > 0.

### Sputnik model 1a: integration into the cell genome

In contrast to Mavirus, Sputnik enters the cell by forming a complex with its host virus (Fig 10). In this model, naïve cells (**C**_**x**_) can be infected by a virus (**G**) or virus-virophage complex (**GV**). Cells infected by the virus (**C**_**g**_) will die at a certain rate and produce viral progeny. Alternatively, cells may be infected by a complex (**C**_**gv**_), release virophages and viral progeny at a level determined by the inhibition parameter (**f**). Since virophages can only enter the cell as a complex, the only way for a virophage to integrate into the cell genome is as a result of an abortive infection. Thus, the integration rate ***λ*** is a compound rate of integration and abortive infection. Cells with a virophage integration (**C**_**i**_), can be infected by a virus (leading to reactivation but no inhibition), or by a virus-virophage complex (leading to inhibition).

**Fig 10.**
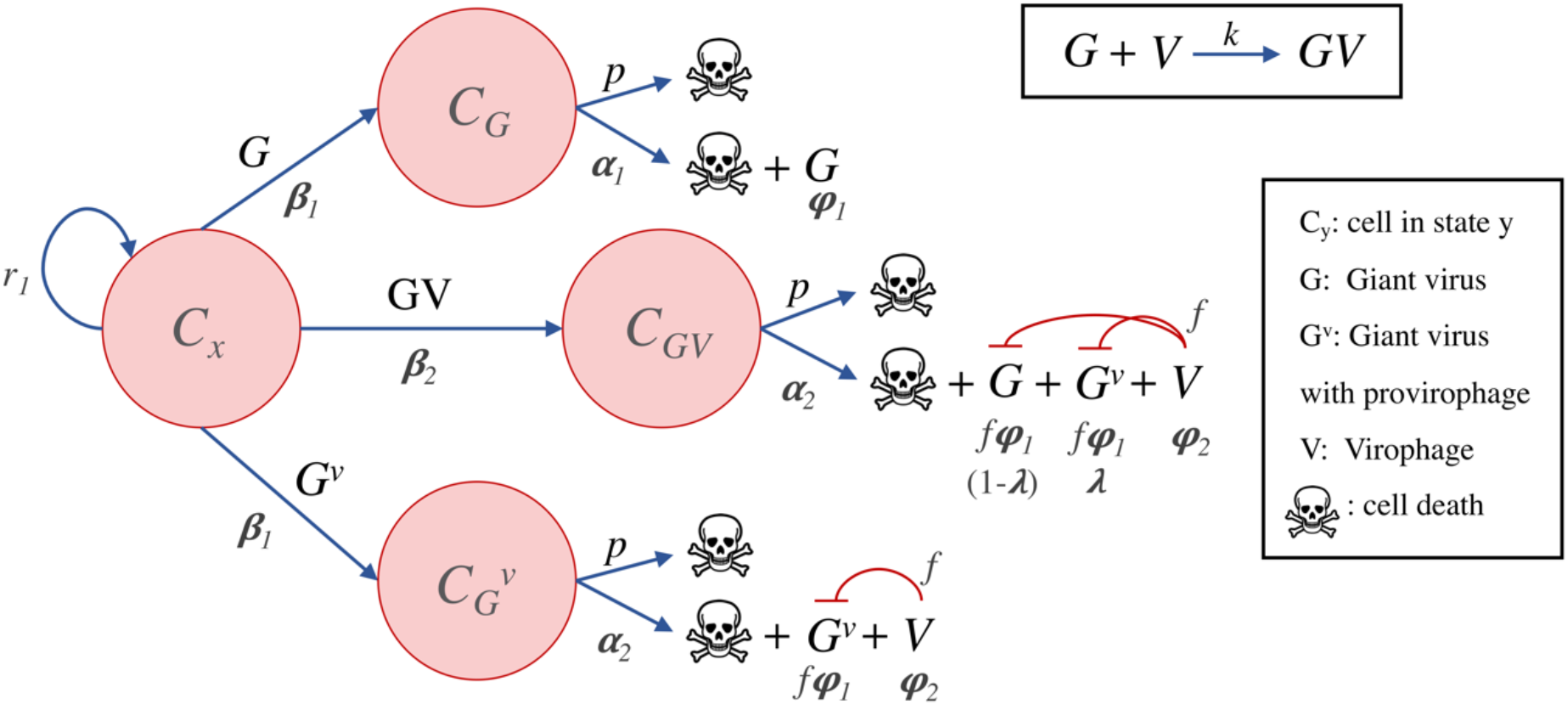
Model of virophage integration into the host virus genome following the Sputnik mechanism. Cells may be infected by a virus (**G**), a virus-virophage complex (**GV**) or a virus carrying a provirophage (**G**_**v**_). During coinfection by a complex (**GV**), a proportion ***λ*\*** of the virus progeny incorporates a virophage into their genomes. Alternatively, when a cell is infected by a virus carrying a provirophage, the virus will produce progeny that also carry the integration, and at the same time will produce expression of virophages.

The Sputnik model with integration into the cell genome is described by a system of 9 coupled ODEs and 15 parameters:

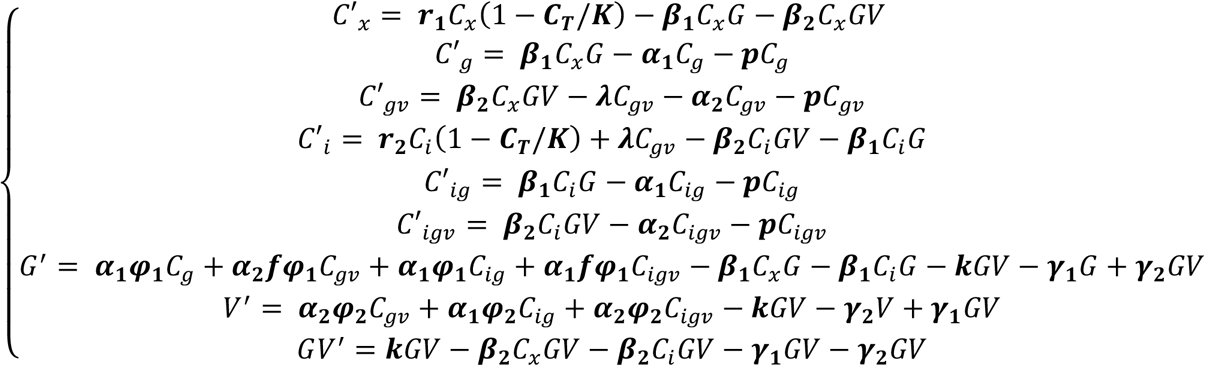

The parameters are the same as outlined for the Mavirus model, except for **β**_**2**_ which refers to the infection rate of the complex (assumed to be equal to **β**_**1**_), and **k** which is the rate of complex formation.

### Sputnik model 1b: integration into the virus genome

We examined a model in which Sputnik can integrate into the virus genome, given that Sputnik provirophages have been found in the genome of *Acanthamoeba polyphaga* Lentille virus (19). In this model, naïve cells can be infected by a virus (**G**), by a complex (**GV**) or a virus with an integrated virophage (**G**_**v**_). Infection by a virus carrying a provirophage results in production of the virus of the same genotype (**G**_**v**_) according to the inhibition parameter (**f**), and release of virophages. Alternatively, infection of a cell by a complex result in production of viral progeny according to **f**, release of virophages and a proportion of viruses with an integrated virophage (the proportion is ***λ*\***). A virus may also infect a naïve cell and replicate without inhibition.

The Sputnik model of integration into the virus genome is described by a system of 8 coupled ODEs and 14 parameters:

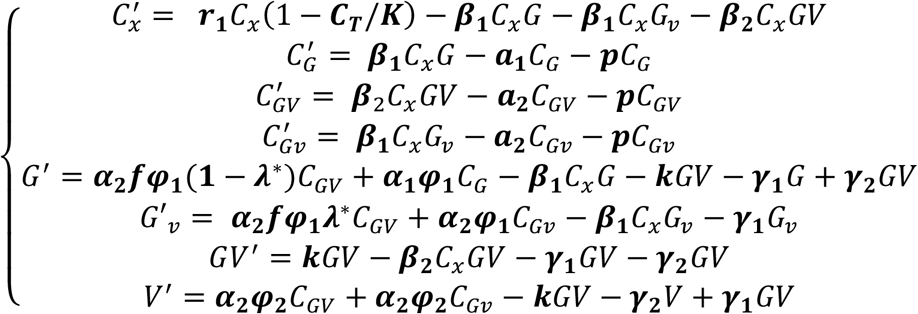

The parameters have the same meanings as those we indicate for the model of Sputnik integration into the cell genome, except for ***λ*\*** which is the proportion of viruses with an integrated virophage that are produced during infection with a complex.

### Numerical integration and choice of parameters

We integrated the models numerically in MATLAB 2020a using the ode45 solver for non-stiff differential equations (45,46). We simulated the models under different scenarios by varying the integration rate of virophages (**λ**, range: [10^−16^, 10^−4^]) and the degree of virophage inhibition (**f**, range: [0 = total inhibition, 1 = no inhibition]). The values for the cell carrying capacity (**K**), cell intrinsic growth rate (**r**), virus burst size (**φ**_**1**_), virophage burst size (**φ**_**2**_) and the rate of complex formation (**k**) have been measured experimentally or estimated from biophysical first principles (47). Therefore, we set our parameters to reflect the order of magnitude of these measurements. Cells carrying a provirophage incurred in a 20% cost of the intrinsic growth rate of naïve cells (**r**_**2**_ = 0.8**r**_**1**_). Our choices for the infection rates of viruses (**β**_**1**_) and virophages (**β**_**2**_), also comprised values in the order of magnitude of those that have been previously used to model this system (32,47). We adjusted the death rate of hosts infected by a virus (**α**_**1**_) or by a virus and virophage (**α**_**2**_), and the decay rates of viruses (**β**_**1**_) and virophages (**β**_**2**_) so that they led to predator-prey oscillations in the presence of neutral virophages; this is our null hypothesis. We used this setup as a baseline scenario to examine the effect of varying the virophage integration rates (***λ***) and degrees of inhibition (**f**) on the system dynamics, which allowed comparison across the different models.

### Agent-based model

We developed an ABM in Julia (48), to study the effect of multicellularity, programmed cell-death and virophage inhibition on the dynamics of the system. Cell, virus and virophage agents were modelled as mutable structures in Julia, with interactions in 3D-space. Cells changed their internal states as they interacted with viruses and virophages (analogous to ODE compartments). Viruses were assumed to follow an independent entry mechanism as in Mavirus. We did not consider dividing cells in this case since this considerably reduces the complexity of the simulations. Therefore, the model follows a single epidemic wave in a population of susceptible hosts.

We started the simulation with 1024 cells, 512 viruses and 2048 virophages. Each agent was assigned a random position in space by sampling a uniform distribution. During the simulation, each agent underwent Gaussian random walks. The size of the displacement for a particle depends on its diffusion coefficient, which according to the Einstein-Stokes equation is a function of the particle size (49). Since virophages are the smallest particles in the system, we set an arbitrary displacement of 1 cell diameter per time step. The displacements of cells and viruses were expressed as a function of the virophage displacement (see S1 Text for derivation, Supporting information).

After every time step, we calculated the distances between cells and viruses, as well as the distance between cells and virophages. If the distance to a cell was less than 1 cell radius, we considered the cell to be infected. Cells infected by viruses died and released viral progeny after the incubation period. If the cell was infected by a virus and a virophage, the cell still died but released virophages and viral progeny according to the inhibition parameter. A cell only infected by a virophage could clear it after a certain number of time steps or alternatively, the virophage could integrate into the cell genome. A virus that infected a cell with an integrated virophage, killed the cell but released virophages and viruses without inhibition.

Multicellular organisms were modelled as cells that were attached to each other in a specified geometry and used the same random displacement vectors. We analysed the dynamics of systems with single-cell, 2-celled, 4-celled, 8-celled and 16-celled agents. Virus induced cell-death was modelled by a constant probability of death per time step, given that a cell is infected by a virus. We analysed three treatments for each population of cell agents: neutral virophages, inhibitory virophages and PCD.

We ran stochastic simulations in 100 replicates for each cell number group/treatment and analysed the effects of treatments and multicellularity by fitting Generalised Additive Models (GAMs). We decided to use GAMs since we observed non-linear relationships in the data and these models allowed testing for multiple effects using smooth functions. Fitting was done using the mgcv package in R (50,51).

## Acknowledgements

We would like to thank Alex Zarebski, Mahan Ghafari and Alan Garfinkel for their helpful comments during preparation of this manuscript. The authors would like to acknowledge use of the University of Oxford Advanced Research Computing facility (http://doi.org/10.5281/zenodo.22558), for carrying out part of this work.

## Supporting information

**S1 Text. Derivation of the displacements of cells and viruses relative to virophages**. These results were used to set the displacements per time step in the ABM simulations.

The Mean Squared Displacement (MSD) is defined as the deviation in the position of a particle with respect to a reference position with time. In statistical mechanics, it is expressed as an ensemble average:

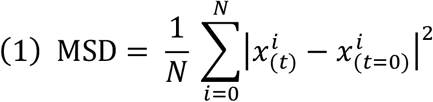

Since we will consider a single particle moving from the origin, we can set N = 1 and *x*^*i*^ at time zero equal to zero. Thus, we have the following expression for the MSD:

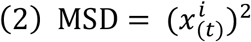

For a random walk in *n* dimensions the MSD is equal to:

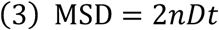

where *D* is the diffusion coefficient for the particle and *t* is time. Since the particles are moving in 3 spatial-dimensions we then have:

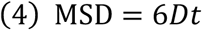

Now we use the Einstein-Stokes equation for the diffusion coefficient of spherical particles at low Reynolds numbers:

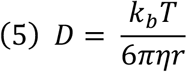

where k_b_ is Boltzmann’s constant, T is the temperature in Kelvin, η is the viscosity of the fluid and r is the radius of a suspended particle.

To relate the displacements to the radius of the particle we substitute (5) in (4), giving:

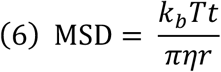

To find the mean displacement we solve for *x* in equation number (2):

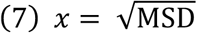

Now we are interested in finding the mean displacement of the virus with respect to the virophage:

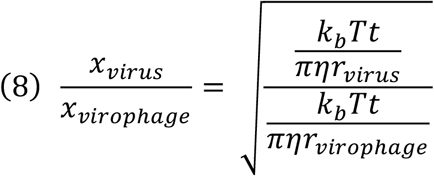

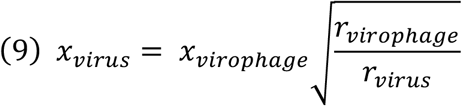

Similarly, the magnitude of the mean displacement of a cell relative to the virophage will be:

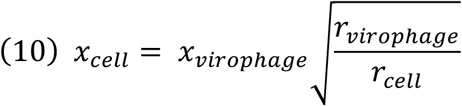

**S1 Fig.**
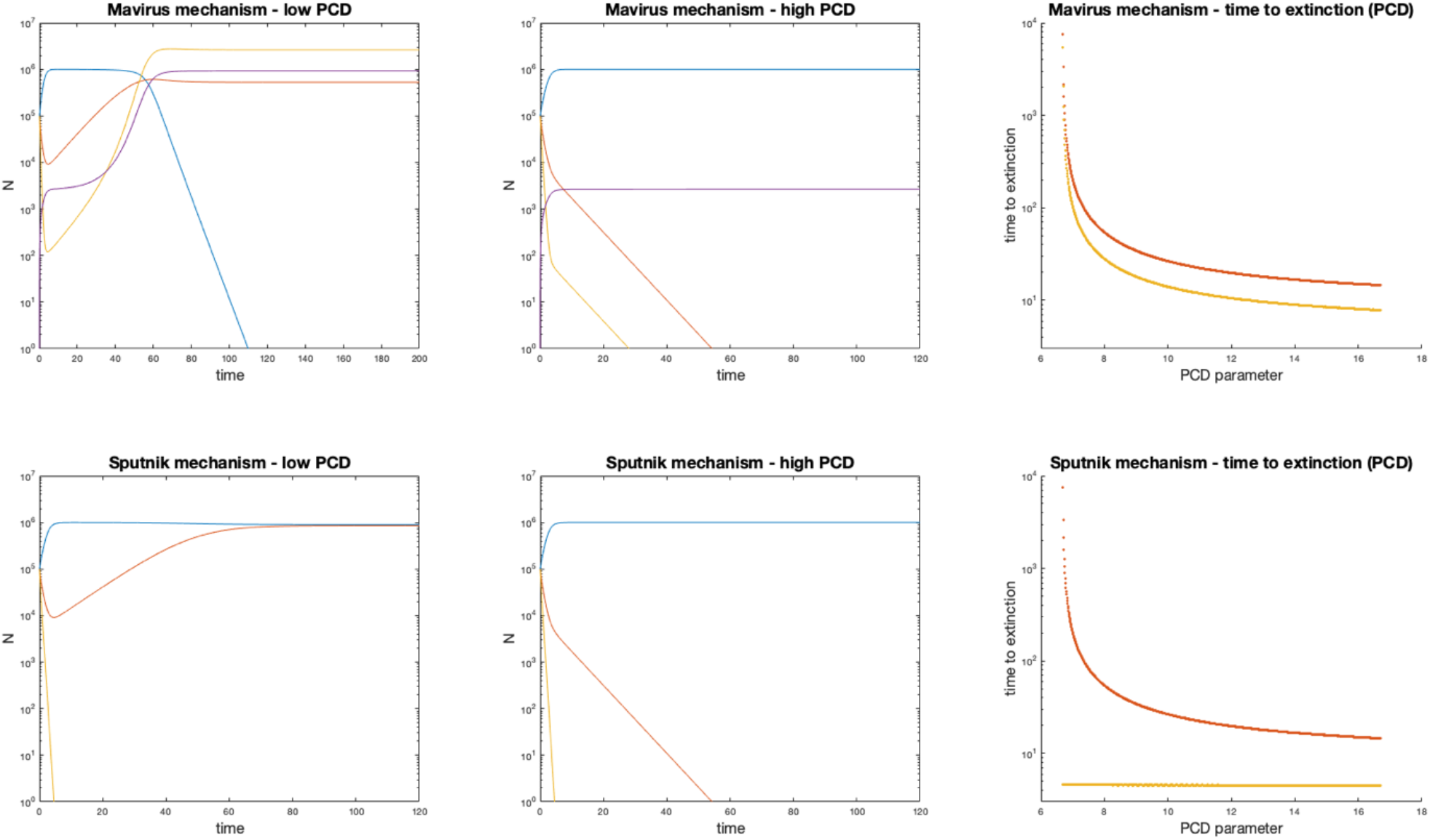
Dynamics of the Mavirus and Sputnik mechanisms in the presence of programmed cell-death (virophages assumed to be neutral). In the time-course diagrams, the blue line represents the population of naïve cells, the purple line represents cells with an integrated virophage, the orange represents viruses and the yellow line represents virophages. **First column:** for the Mavirus model and a low PCD parameter (p = 6), we observe fixation of cells with the integrated virophage and stable coexistence of viruses and neutral virophages. In the Sputnik model, virophages go extinct and naïve cells persist in the population. **Second column:** increasing the PCD parameter (p = 8) leads to extinction of viruses and virophages in both cases. An integration polymorphism is observed in the Mavirus model, while naïve cells grow to carrying capacity in the Sputnik model. **Third column:** transition to the extinction regime occurs with a PCD parameter greater than 6.6. Time to virus extinction decreases as a function of the PCD parameter in both models, and for virophages in the Mavirus model. Parameters for model 1: **α**_**1**_ = 1, **α**_**2**_ = 0.9, **β**_**1**_ = 10^−7^, **β**_**2**_ = 10^−6^, **γ**_**1**_ = 1.2, **γ**_**2**_ = 2.6, ***λ*** = 10^−7^, **φ**_**1**_ = 100, **φ**_**2**_ = 1000, **f** = 1, **K** = 10^6^, **r**_**1**_ = 1, **r**_**2**_ = 0.8. Parameters for model 2 are the same except for _**2**_ = 10^−7^ and **k** = 8·10^−7^. Initial conditions: **C**_**x**,**0**_ = 10^5^, **G**_**0**_ = 10^5^, **V**_**0**_ = 10^5^.

**S2 Fig.**
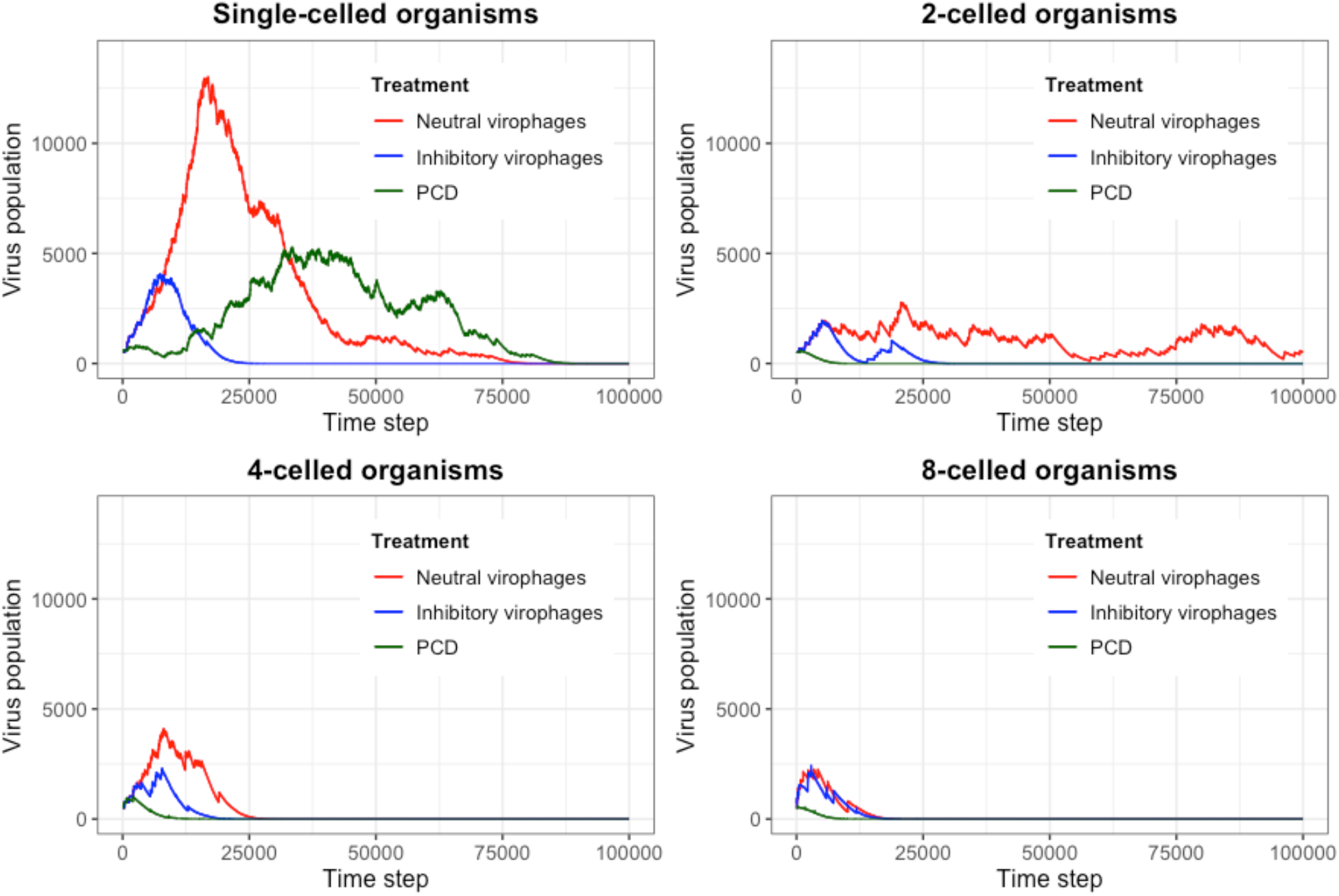
Time-course of the population of viruses during an ABM simulation. The total virus population during a stochastic simulation is shown as a function of time and grouped by the number of cells per organism and treatment. The highest virus wave is observed in the population of single-celled organisms in the presence of neutral virophages. By comparison, treatment with inhibitory virophages and PCD show lower maxima of the virus waves. The effect of multicellularity can also be observed in the lower virus waves from the single-celled to 8-celled cases. Random seed = 1234.

**S3 Fig.**
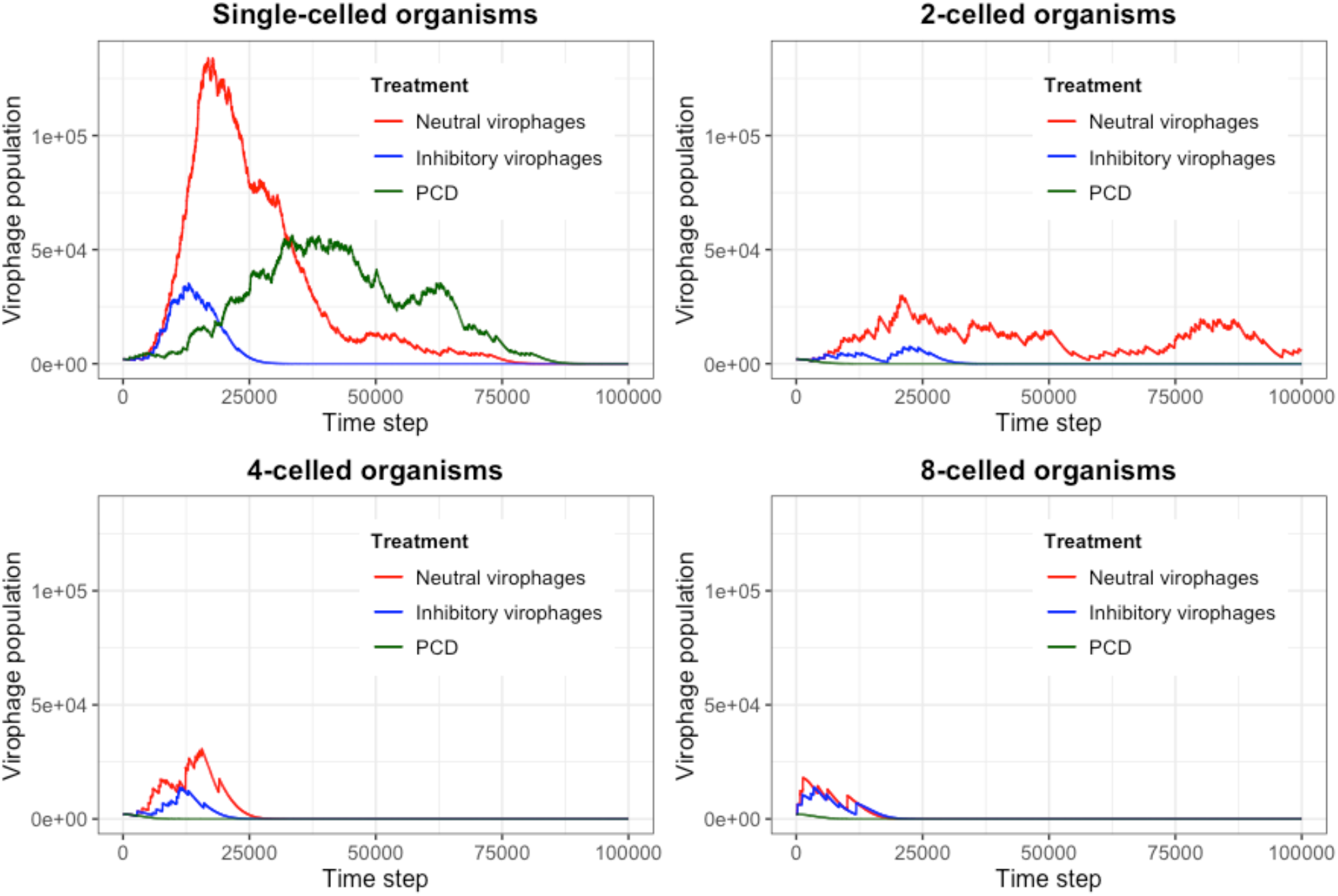
Time-course of the population of virophages during an ABM simulation. The total virophage population during a stochastic simulation is plotted as a function of time and grouped by the number of cells per organism and treatment. We observe the same pattern as for the virus population, where the highest virophage numbers were attained in the simulation with neutral virophages and lower for inhibitory virophages and PCD. In contrast to viruses, virophages attain much larger population sizes. Multicellularity also has an effect on reducing the maximum size of the virophage waves. Random seed = 1234.

**S4 Fig.**
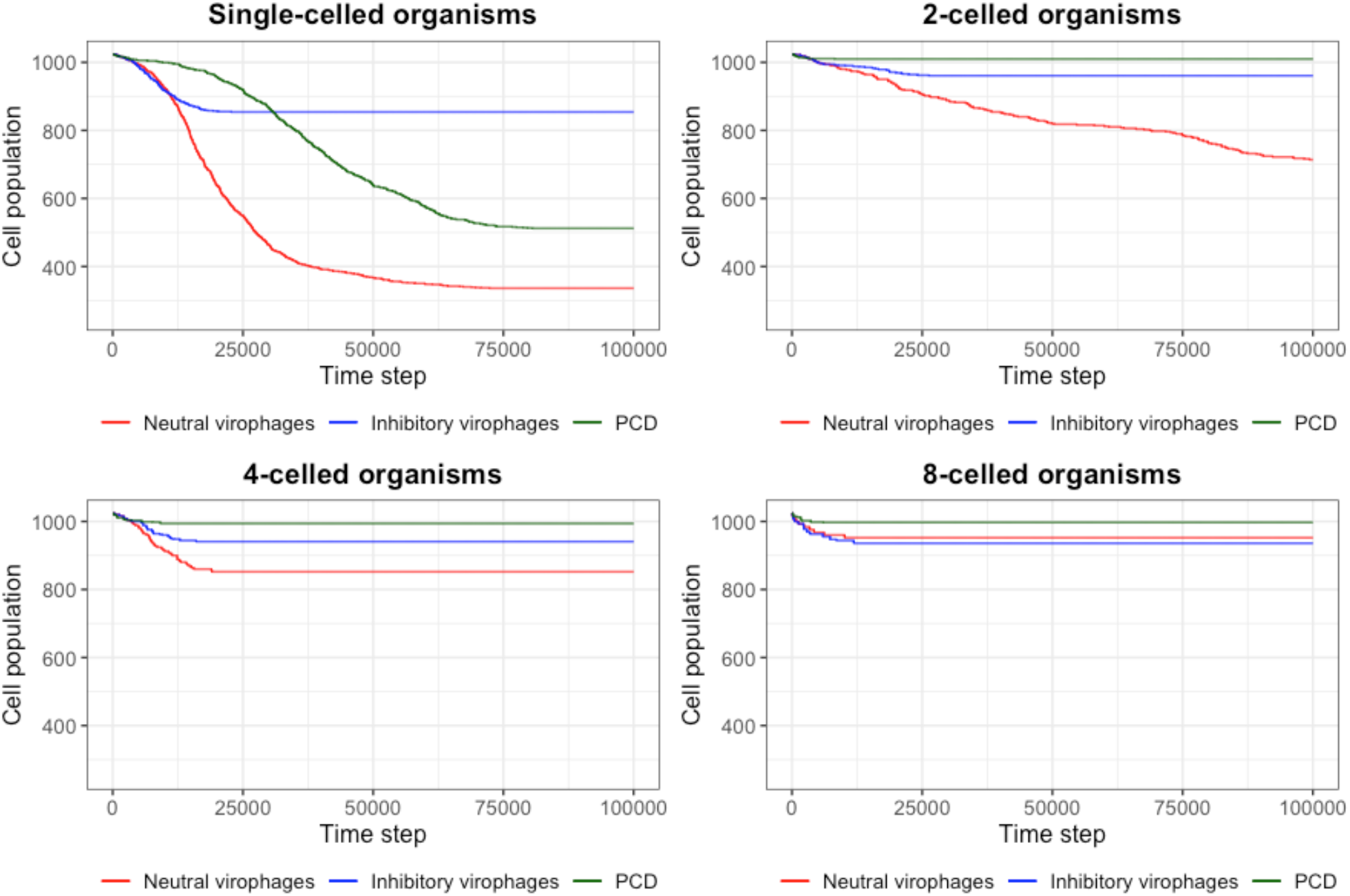
Time-course of the number of surviving cells during an ABM simulation. The total number of cells during a stochastic simulation is shown as a function of time and grouped by the number of cells per organism and treatment. We can observe a greater survival of cells in treatments with PCD and inhibitory virophages, while the lowest survival occurred in the treatments with neutral virophages. Grouping cells together also provided an advantage since the survival of cells increases with increasing number of cells per organism. Random seed = 1234.

**S1 Video. Simulation of a population of single-celled organisms interacting with viruses and neutral virophages**. The simulation was started with 1024 cells, 512 viruses and 2048 virophages. Cells are represented as blue, viruses are red and virophages are green. The axes are in micrometres. Random seed = 1234.

**S2 Video. Simulation of a population of single-celled organisms interacting with viruses and inhibitory virophages**. The simulation was started with 1024 cells, 512 viruses and 2048 virophages. Cells are represented as blue, viruses are red and virophages are green. The axes are in micrometres. Random seed = 1234.

**S3 Video. Simulation of a population of single-celled organisms interacting with viruses and neutral virophages, which can undergo virus-induced PCD**. The simulation was started with 1024 cells, 512 viruses and 2048 virophages. Cells are represented as blue, viruses are red and virophages are green. The axes are in micrometres. Random seed = 1234.

**S4 Video. Simulation of a population of 2-celled organisms interacting with viruses and neutral virophages**. The simulation was started with 1024 cells, 512 viruses and 2048 virophages. Cells are represented as blue, viruses are red and virophages are green. The axes are in micrometres. Random seed = 1234.

**S5 Video. Simulation of a population of 2-celled organisms interacting with viruses and inhibitory virophages**. The simulation was started with 1024 cells, 512 viruses and 2048 virophages. Cells are represented as blue, viruses are red and virophages are green. The axes are in micrometres. Random seed = 1234.

**S6 Video. Simulation of a population of 2-celled organisms interacting with viruses and neutral virophages, which can undergo virus-induced PCD**. The simulation was started with 1024 cells, 512 viruses and 2048 virophages. Cells are represented as blue, viruses are red and virophages are green. The axes are in micrometres. Random seed = 1234.

**S7 Video. Simulation of a population of 4-celled organisms interacting with viruses and neutral virophages**. The simulation was started with 1024 cells, 512 viruses and 2048 virophages. Cells are represented as blue, viruses are red and virophages are green. The axes are in micrometres. Random seed = 1234.

**S8 Video. Simulation of a population of 4-celled organisms interacting with viruses and inhibitory virophages**. The simulation was started with 1024 cells, 512 viruses and 2048 virophages. Cells are represented as blue, viruses are red and virophages are green. The axes are in micrometres. Random seed = 1234.

**S9 Video. Simulation of a population of 4-celled organisms interacting with viruses and neutral virophages, which can undergo virus-induced PCD**. The simulation was started with 1024 cells, 512 viruses and 2048 virophages. Cells are represented as blue, viruses are red and virophages are green. The axes are in micrometres. Random seed = 1234.

**S10 Video. Simulation of a population of 8-celled organisms interacting with viruses and neutral virophages**. The simulation was started with 1024 cells, 512 viruses and 2048 virophages. Cells are represented as blue, viruses are red and virophages are green. The axes are in micrometres. Random seed = 1234.

**S11 Video. Simulation of a population of 8-celled organisms interacting with viruses and inhibitory virophages**. The simulation was started with 1024 cells, 512 viruses and 2048 virophages. Cells are represented as blue, viruses are red and virophages are green. The axes are in micrometres. Random seed = 1234.

**S12 Video. Simulation of a population of 8-celled organisms interacting with viruses and neutral virophages, which can undergo virus-induced PCD**. The simulation was started with 1024 cells, 512 viruses and 2048 virophages. Cells are represented as blue, viruses are red and virophages are green. The axes are in micrometres. Random seed = 1234.

**S13 Video. Simulation of a population of 16-celled organisms interacting with viruses and neutral virophages**. The simulation was started with 1024 cells, 512 viruses and 2048 virophages. Cells are represented as blue, viruses are red and virophages are green. The axes are in micrometres. Random seed = 1234.

**S14 Video. Simulation of a population of 16-celled organisms interacting with viruses and inhibitory virophages**. The simulation was started with 1024 cells, 512 viruses and 2048 virophages. Cells are represented as blue, viruses are red and virophages are green. The axes are in micrometres. Random seed = 1234.

**S15 Video. Simulation of a population of 16-celled organisms interacting with viruses and neutral virophages, which can undergo virus-induced PCD**. The simulation was started with 1024 cells, 512 viruses and 2048 virophages. Cells are represented as blue, viruses are red and virophages are green. The axes are in micrometres. Random seed = 1234.

